# MicroRNA-29a-5p attenuates hemorrhagic transformation and improves outcomes after mechanical reperfusion for acute ischemic stroke

**DOI:** 10.1101/2024.02.13.580104

**Authors:** Chang-Luo Li, Jin-Kun Zhuang, Zhong Liu, Zhong-Run Huang, Chun Xiang, Qian-Yu Chen, Ze-Xin Chen, Zhong-Song Shi

**Affiliations:** Department of Neurosurgery (C-LL, J-KZ, Z-RH, CX, Z-SS), Sun Yat-sen Memorial Hospital, Sun Yat-sen University, Guangzhou, China; RNA Biomedical Institute (C-LL, J-KZ, ZL, Z-RH, Q-YC, Z-XC, Z-SS), Sun Yat-sen Memorial Hospital, Sun Yat-sen University, Guangzhou, China; Nanhai Translational Innovation Center of Precision Immunology (C-LL, J-KZ, Z-RH, Q-YC, Z-XC, Z-SS), Sun Yat-sen Memorial Hospital, Sun Yat-sen University, Foshan, China; Department of Neurosurgery (ZL), Zhongshan Hospital of Xiamen University, School of Medicine, Xiamen University, Xiamen, China; Guangdong Province Key Laboratory of Brain Function and Disease (Z-SS), Sun Yat-sen University, Guangzhou, China

**Keywords:** Acute ischemic stroke, Hemorrhagic transformation, Cerebral ischemia-reperfusion injury, Oxygen-glucose deprivation reoxygenation, MicroRNA, Astrocyte

## Abstract

**Background:** Hemorrhage transformation (HT) following endovascular reperfusion treatment is associated with worse clinical outcomes in acute ischemic stroke patients. MicroRNA (miR) modulates several aspects of cerebral ischemia-reperfusion injury, including blood-brain barrier (BBB) integrity, inflammation, oxidative stress, and apoptosis, significantly impacting cerebral recovery and function. This study investigated the role of astrocytic miR-29a-5p in HT in the transient middle cerebral artery occlusion (MCAO) model and oxygen-glucose deprivation reoxygenation (OGD/R) model of astrocytes.

**Methods:** MiR-29a-5p expression in the OGD/R astrocyte model was assessed. The astrocyte injury, the expression of A1 and A2 phenotypes of reactive astrocytes, and the regulation of miR-29a-5p target genes were evaluated after the miR-29a-5p intervention. A mechanical reperfusion-induced HT model was established in hyperglycemic rats using 5-hour MCAO following reperfusion at 6 hours. MiR-29a-5p agomir was administered intravenously before reperfusion. Infarct volume, HT, BBB damage, neurological score, the expression of miR-29a-5p, and its target genes were evaluated.

**Results:** MiR-29a-5p expression decreased in OGD/R-treated astrocytes and the peri-infarction tissue and blood of the MCAO model. Elevating miR-29a-5p levels reduced astrocyte injury, suppressed neurotoxic A1 astrocyte markers (C3, Fkbp5, and Serping1), while enhanced neuroprotective A2 astrocyte markers (S100a10 and Emp1) in the OGD/R and MCAO models. Intravenous administration of miR-29a-5p agomir increased the expression of miR-29a-5p and reduced infarct volume, reperfusion-induced HT, and BBB breakdown after ischemia, improving neurological outcomes in the MCAO model. Overexpression of miR-29a-5p effectively suppressed the expression of its direct target genes, glycogen synthase kinase 3 beta and aquaporin 4 in the OGD/R and MCAO models.

**Conclusions:** MiR-29a-5p alleviates astrocyte injury and regulates A1 and A2 astrocyte markers, glycogen synthase kinase 3 beta, and aquaporin 4 in astrocytes subjected to ischemia-reperfusion injury. Astrocytic miR-29a-5p may be a protective target for reducing HT and improving outcomes following mechanical reperfusion in acute ischemic stroke.

## 1. Introduction

Acute ischemic stroke (AIS) patients with intracranial large-vessel occlusion can benefit from earlier treatment with endovascular thrombectomy and intravenous alteplase [1–3]. However, the cerebral ischemia-reperfusion injury that often follows reperfusion treatments introduces new challenges, including hemorrhagic transformation (HT) and exacerbated inflammatory responses, which are detrimental to recovery and are predictive of poor long-term outcomes in AIS patients [4–8]. Emerging research has highlighted the pivotal role of non-coding RNAs in these processes. MicroRNA (miR) modulates several aspects of cerebral ischemia-reperfusion injury, including blood-brain barrier (BBB) integrity, inflammation, oxidative stress, and apoptosis, significantly impacting cerebral recovery and function [9–14].

The miR-29 family, including miR-29a, miR-29b, and miR-29c, is significantly downregulated in cerebral ischemia-reperfusion injury, which affects cellular responses to ischemic damage [15–18]. Specifically, miR-29a-5p influences neuroglial interactions, impacting microglial activation and glutamate release and affecting the volume of ischemic damage and infarcts seen in animal stroke models [19]. Studies suggest that nicotinamide adenine dinucleotide phosphate oxidase inhibitor, by increasing miR-29a-5p levels, help to reduce cerebral infarct volumes and HT after reperfusion in ischemic rat tissues [20,21]. By regulating these key processes, miR-29a-5p plays a pivotal role in reducing the detrimental effects of ischemia-reperfusion injury and promoting neuroprotection. Non-coding RNAs, including microRNAs, have emerged as important diagnostic and prognostic blood biomarkers for AIS due to their stability in body fluids and specific expression patterns in response to ischemic injury [9,10,14]. MiR-29a-5p was found to be significantly downregulated in the plasma of stroke patients, correlating with the severity of brain injury and clinical outcomes [19]. This makes miR-29a-5p a valuable biomarker for early diagnosis and prognosis, aiding in the prediction of patient outcomes and the personalization of therapeutic strategies.

Astrocytes, the predominant type of glial cells in the central nervous system, play critical roles in supporting neuronal function and contributing to neurodegenerative processes [22]. In ischemic stroke, reactive astrocytes can adopt a neurotoxic A1 phenotype or a neuroprotective A2 phenotype. A1 astrocytes exacerbate neuronal damage by releasing pro-inflammatory cytokines and reactive oxygen species, while A2 astrocytes promote recovery by secreting neurotrophic factors and anti-inflammatory cytokines [23,24].

MiR-29a is highly expressed in astrocytes. Despite its protective roles, the function of miR-29a-5p in HT remains inadequately understood. This study aims to clarify the mechanisms by which astrocytic miR-29a-5p regulates HT using the oxygen-glucose deprivation and reoxygenation (OGD/R) model and a mechanical reperfusion-induced HT rat model. The focus is on how miR-29a-5p influences the expression between neurotoxic A1 astrocytes, which can exacerbate injury, and neuroprotective A2 astrocytes, which aid recovery. By understanding the interactions between miR-29a-5p and its target genes within astrocytes, this study aims to uncover novel molecular mechanisms that could be targeted to reduce HT following endovascular reperfusion treatment, offering the potential of miR-29a-5p as a therapeutic target in stroke treatment.

## 2. Materials and Methods

### 2.1. Establishing a mechanical reperfusion-induced HT model

The experimental protocol received approval from our institution’s Institutional Animal Care and Use Committee. Adult male Sprague-Dawley rats (8-10 weeks, 250-280g) were utilized to establish a hyperglycemia-associated reperfusion-induced HT model following established protocols [20,21,25,26]. The intraluminal filament technique was performed to have the left middle cerebral artery occlusion (MCAO) for 5 hours, followed by reperfusion at 6 hours. Anesthesia was initiated with 5% isoflurane in a 70% nitrogen/30% oxygen mixture and maintained with 2% isoflurane via a facemask. Body temperature was stabilized at 37.0°C using a heating pad. Immediate reperfusion was achieved by filament withdrawal after the ischemic period. Animals were euthanized 6 hours post-reperfusion (11 hours post-ischemia onset) for tissue collection.

Acute hyperglycemia was induced through intraperitoneal administration of 50% dextrose (6ml/kg) 15 minutes pre-occlusion, maintaining blood glucose >16.7 mmol/L. Sustained hyperglycemia during ischemia was achieved via supplemental 50% dextrose injections (1.5ml/kg) at 1, 2, 3, and 4 hours post-MCAO. Blood glucose monitoring occurred via tail vein sampling at baseline and 0.5, 1, 2, and 4 hours post-dextrose administration. The hyperglycemic condition increases the extent of cerebral hemorrhage after ischemia-reperfusion in the MCAO model.

### 2.2. MiR-29a-5p agomir administration in HT model

In the study, 151 rats met the criteria of successful surgery and acute hyperglycemia. One rat was excluded because of subarachnoid hemorrhage after the surgery. First, 30 hyperglycemic rats were used to assess the dynamic analysis of the levels of miR-29a-5p in the peri-infarction tissue and whole blood. Rats were randomly enrolled into five groups: sham group, 5-hour MCAO following reperfusion at 0, 1, 3, and 6 hours (6 rats per group).

Then, rats were used to assess the role of miR-29a-5p in HT and the underlying mechanisms. One hundred and twenty hyperglycemic rats were randomly assigned into four groups: (1) sham-operated, (2) ischemia-reperfusion, (3) ischemia-reperfusion and microRNA control, (4) ischemia-reperfusion and miR-29a-5p agomir group (30 rats per group). Rats received an intravenous injection of miR-29a-5p agomir or microRNA control (Gene Pharma) at a dose of 50 nmol/kg body weight 30 minutes before reperfusion. Rats were sacrificed 6 hours after reperfusion, and brains were harvested for the measurements.

Ischemic hemisphere peri-infarct regions were dissected using previously described methods [27]. Specifically, after delineating the midline separating the ischemic and non-ischemic hemispheres, a longitudinal incision was made along the sagittal plane, 2 mm lateral to the midline, extending from the top to the bottom across each hemisphere. Subsequently, an oblique transverse incision was made to divide the brain tissue into the infarction core (striatum and upper cerebral cortex) and the peri-infarct region (adjacent cerebral cortex). The peri-infarct tissue was used for further RT-PCR and Western blot study. The whole blood was collected and stored in the PAXgene Blood RNA tube (PreAnalytiX GmbH) to extract the total RNA.

### 2.3. Neurological evaluation

Neurological assessments were conducted pre-ischemia and 6 hours post-reperfusion using a 5-point scale (0 = no neurological deficits; 1 = failure to extend contralateral forepaw fully; 2 = reduced resistance to lateral push; 3 = spontaneous circling to contralateral side; 4 = absence of spontaneous movement or unconsciousness)

### 2.4. Cerebral infarction quantification

Infarct volumes were determined via 2,3,5-triphenyl tetrazolium chloride (TTC) staining 11 hours post-ischemia. Six 2-mm coronal sections were incubated with 2% TTC solution (Sigma) for 20 minutes at room temperature. Edema-corrected infarct volume was calculated using ImageJ software as the following formula: (contralateral hemisphere volume - non infarct ipsilateral hemisphere volume)/contralateral hemisphere volume.

### 2.5. BBB integrity assessment

Evans blue extravasation was evaluated 11 hours after ischemia. Evans blue solution (Sigma) was administered via the tail vein 8 hours after ischemia. Three hours later, vascular clearance via cardiac perfusion preceded brain dissection. Hemispheric samples were homogenized in trichloroacetic acid, centrifuged (10,000g, 30 minutes), and spectrophotometrically analyzed at 620nm. Evans blue extravasation index was expressed as the ratio of absorbance intensity in the ischemic hemisphere to that in the non-ischemic hemisphere.

### 2.6. Measurement of cerebral hemorrhage

Cerebral hemoglobin content was measured spectrophotometrically 11 hours post-ischemia. Homogenized hemispheric samples in PBS were centrifuged (13,000g, 30 minutes), mixed with Drabkin’s reagent (Sigma), and analyzed by spectrophotometry at 540nm. The hemoglobin content index was calculated as hemoglobin content in the ipsilateral hemisphere divided by that in the contralateral hemisphere.

### 2.7. Astrocyte cell culture and OGD/R model

Primary astrocytes were isolated from neonatal Sprague-Dawley rat cortices as previously reported [28]. The cortex was isolated and then digested with 0.25% trypsin (Gibco, 25200072) at 37 °C for 20 minutes. The digestion was terminated with a complete medium containing DMEM-F12 (Gibco, C11330500B), 10% FBS (Gibco,10099141), and 1% Penicillin-Streptomycin (Gibco, 15140122). The cell suspension was collected using a 70 µM nylon cell filter and centrifuged at 1000 rpm for 5 minutes. Cells were resuspended using the complete medium, plated into 75 cm2 flasks, and then cultured in a humidified incubator with 5% CO2 at 37°C, with the complete medium replaced every 2-3 days. We identified the purity of astrocytes using immunofluorescence and used the third-generation cells to perform the subsequent OGD/R experiments.

Astrocytes underwent OGD/R in hypoxic chambers (Stemcell) (95% N₂ and 5% CO₂, Stemcell) with glucose-free medium (Gibco, 11966025) for 6 hours, followed by normoxic reoxygenation in glucose-containing complete medium. We collected the cells 0, 3, 6, 12, and 24 hours after reoxygenation. Astrocytes did not receive OGD as the control.

### 2.8. Prediction of hub target genes for miR-29a-5p

The predicted target genes of miR-29a-5p were identified using the miRDB and TargetScan databases. We performed Kyoto Encyclopedia of Genes and Genomes (KEGG) pathway and Gene Ontology functional enrichment analyses to identify the biological functions of the predicted target genes using DAVID Bioinformatics Resources [29]. The predicted genes from the top ten most enriched KEGG pathways were used to search for the interactions of the target genes using the STRING tool. We obtained the protein-protein interaction network and investigated the top twenty hub genes using CytoHubba.

### 2.9. Transfection of miR-29a-5p mimics and inhibitors

Astrocytes were transfected with miR-29a-5p mimics and inhibitors, small interfering RNA (siRNA) for the predicted target gene, including glycogen synthase kinase 3 beta (Gsk3b), aquaporin 4 (Aqp4), and FKBP prolyl isomerase 5 (Fkbp5). The optimal concentrations of the miR-29a-5p mimics and miR-29a-5p inhibitors were 50nM and 100nM. The optimal sequence of siRNA for target gene of rat Gsk3b was as follows: sense strand: GGGAGCAAAUUAGAGAAAUTT, antisense strand: AUUUCUCUAAUUUGCUCCCTT. The optimal sequence of siRNA for target gene of rat Aqp4 was as follows: sense strand: GCACACGAAAGAUCAGCAUTT, antisense strand: AUGCUGAUCUUUCGUGUGCTT. The optimal sequence of siRNA for target gene of rat Fkbp5 was as follows: sense strand: GUAUCUUGGACCACAAUAUTT, antisense strand: AUAUUGUGGUCCAAGAUACTT. The miR-29a-5p mimics and inhibitors were synthesized by RiboBio. siRNA genes were synthesized by Gene Pharma.

Astrocytes were performed OGD/R and divided into six groups: blank control, OGD/R, OGD/R receiving negative control microRNA, OGD/R receiving the miR-29a-5p mimics, OGD/R receiving the miR-29a-5p inhibitors, and siRNA gene (Gsk3b, Aqp4 and Fkbp5). Astrocytes were transfected with siRNA using Lipofectamine 3000 (Invitrogen, L3000015). After 48 hours of incubation, OGD/R was performed, and then RT-PCR was performed to evaluate the transfection efficacy.

### 2.10. Cell Counting Kit-8 (CCK-8) assay

A CCK-8 assay was used to detect the viability of cells in the OGD/R astrocyte model. Astrocytes were seeded into 96-well plates (5×10^4^ cells/well) and then transfected with miRNA-29a-5p mimics (50 nM), miRNA-29a-5p inhibitors (100 nM), or negative control miRNA. After 48 hours of incubation, 20 µl of CCK-8 solution (Dojindo Laboratories, CK04, Japan) was added to each well and incubated for 1 hour. The absorbance of each well was measured at 450 nm.

### 2.11. Lactate dehydrogenase (LDH) assay

An LDH Cytotoxicity Colorimetric Assay Kit was used to analyze LDH release in the medium in the OGD/R astrocyte model. Astrocytes were divided into control, OGD/R, OGD/R receiving microRNA control, OGD/R receiving miR-29a-5p mimics, and OGD/R receiving miR-29a-5p inhibitors group. We collected and centrifuged the supernatants of each group. The LDH levels in supernatants were measured using an LDH Cytotoxicity Assay Kit (Beyotime Biotechnology, C0016, China). We measured the absorbance of each group at 490 nm to determine the LDH activity after 30 minutes of incubation.

### 2.12. Quantitative RT-PCR

The total RNA of the astrocytes in the OGD/R model and the peri-infarction tissue and whole blood in the MCAO model were extracted using a Trizol reagent (Invitrogen). The mRNAs of genes from the prediction analysis, A1 reactive astrocyte markers [complement C3 (C3), Fkbp5, serpin family G member 1 (Serping1), glycoprotein alpha-galactosyltransferase 1 (Ggta1), guanylate binding protein 2 (Gbp2)], and A2 reactive astrocyte markers [S100 calcium binding protein A10 (S100a10), pentraxin 3 (Ptx3), sphingosine-1-phosphate receptor 3 (S1pr3), epithelial membrane protein 1 (Emp1), CD109 molecule (Cd109)] were assessed. We used the SYBR green real-time RT-PCR and the TaqMan stem-loop method to assess the level of miR-29 and mRNA [20,21]. Quantitative RT-PCR was performed using the Applied Biosystem PCR System. We normalized the levels of mRNA and miR-29a-5p using β-actin and U6 as internal controls and used the 2-ΔΔCT method to calculate them. Each sample was tested in triplicate. The mRNA and miR-29a-5p primer were designed and synthesized by RiboBio, China. The primer sequences are shown in Table 1.

**Table 1.**
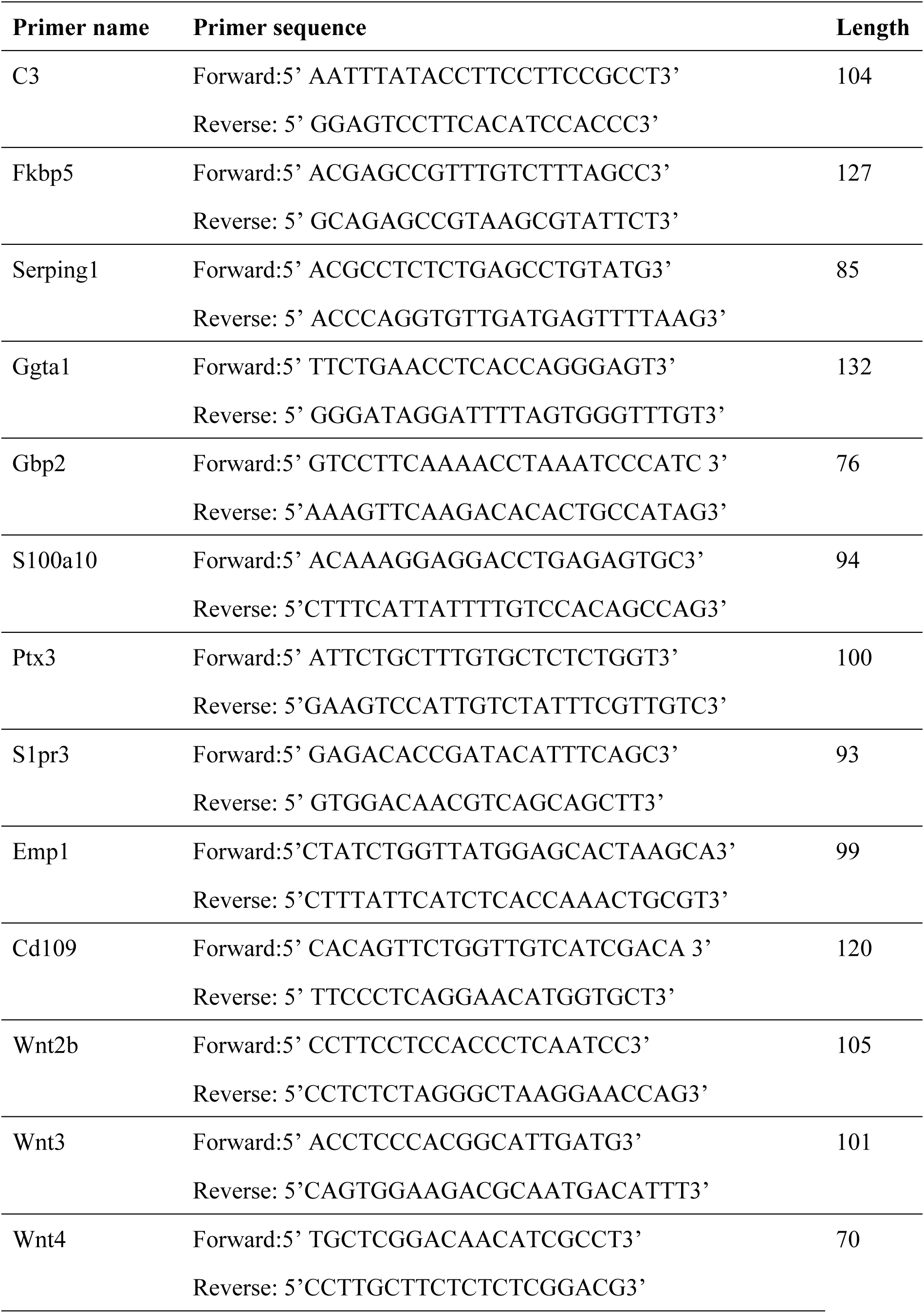
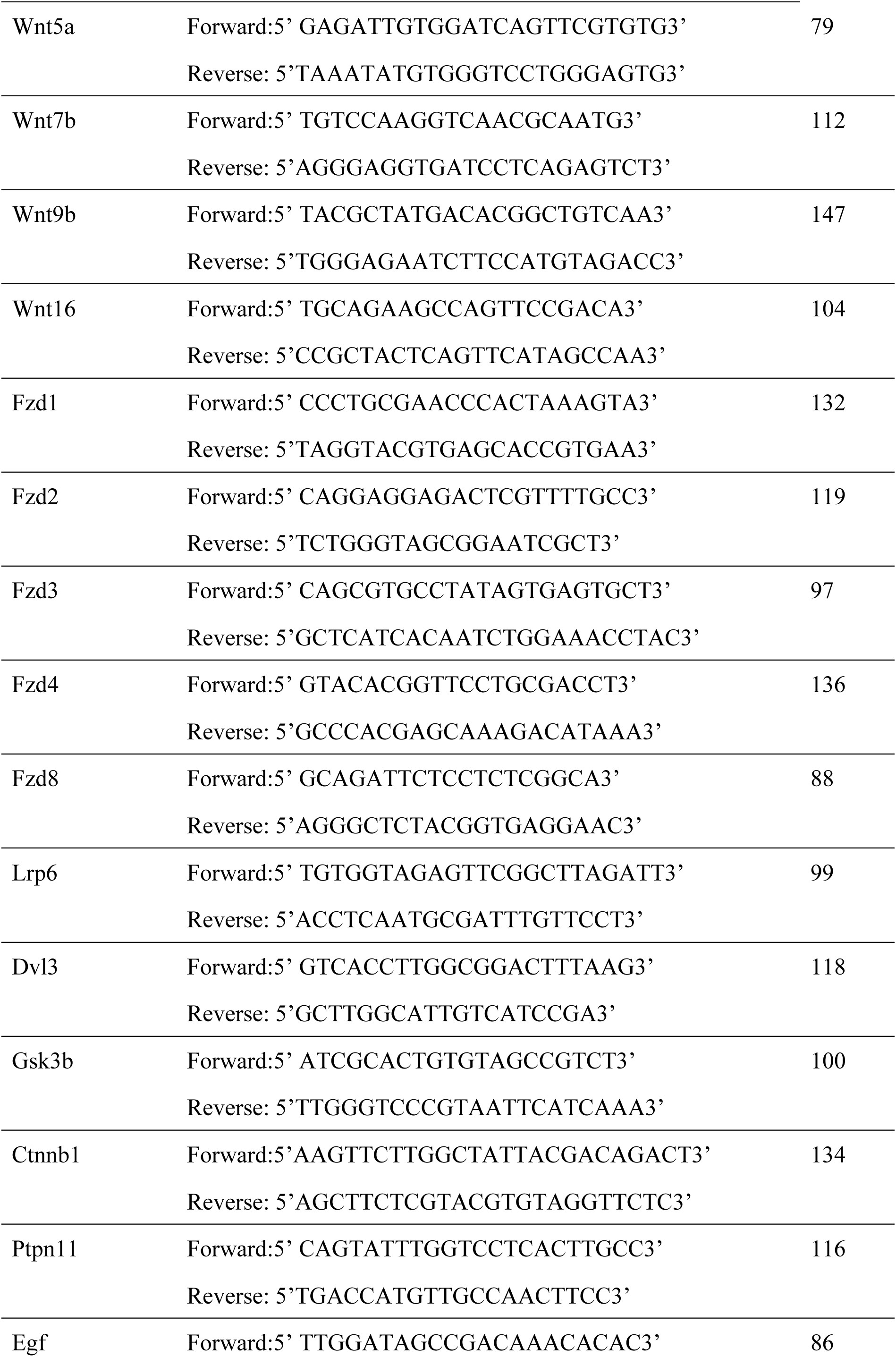
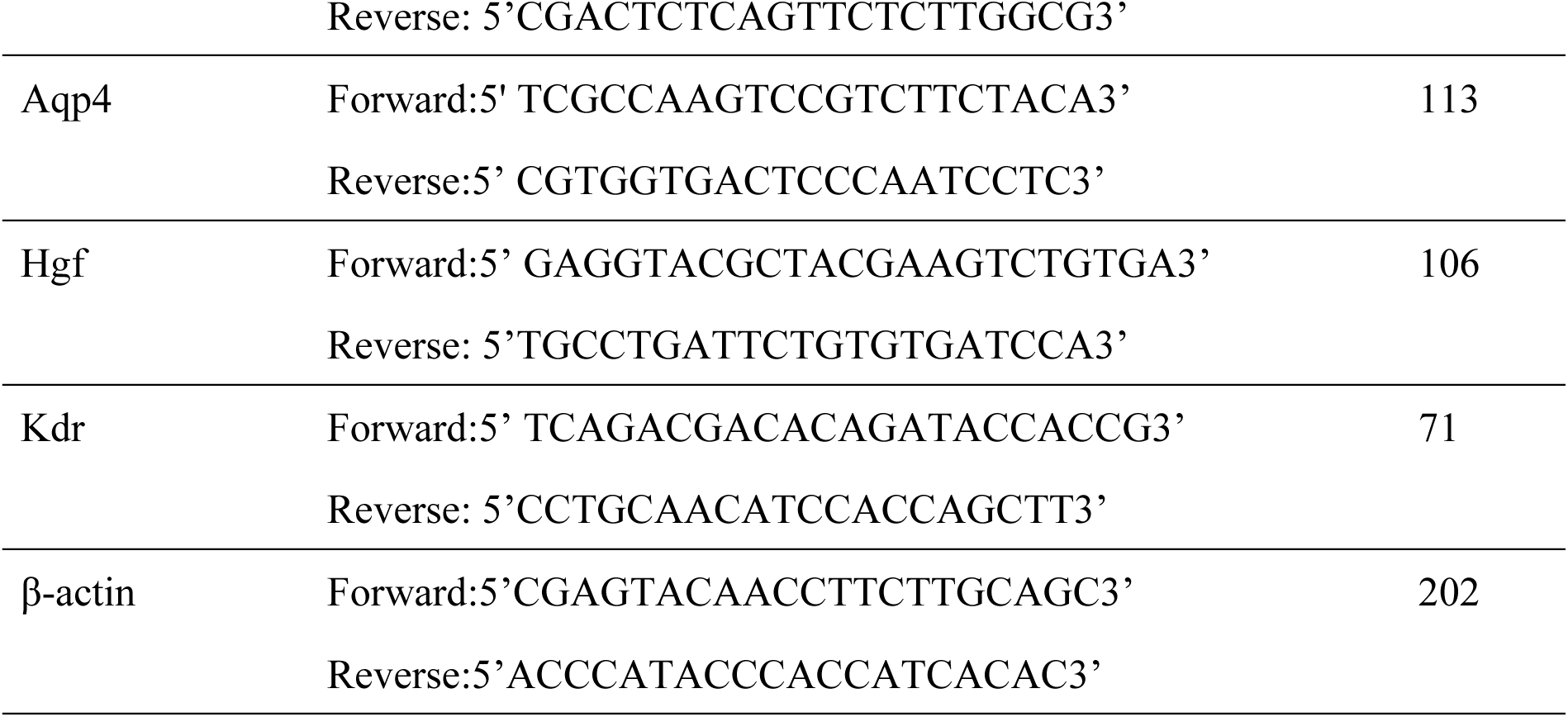
Sequences of the primer for mRNAs used in the qRT-PCR.

### 2.13. Western blot

We extracted the total protein of the astrocytes following OGD/R and the peri-infarction tissue of the MCAO model using a bicinchoninic acid protein assay and assessed the levels of GSK3β, AQP4, and FKBP5. The anti-GSK3β (Abcam, ab131356, 1:1000), anti-AQP4 (Abcam, ab9512, 1:1000), and anti-FKBP5 (Abcam, ab126715, 1:1000) antibodies were visualized using an HRP-conjugated secondary antibody and enhanced chemiluminescence. The expression bands of protein were detected using a molecular imager system. The density of bands was normalized to β-actin (Abcam, ab227387,1:1000). Each sample was tested in triplicate. Image J software was used to quantify the optical density value of proteins.

### 2.14. Dual-luciferase reporter assay

Dual-luciferase activity technology was used to clarify the relationship between miR-29a-5p and Gsk3b and Aqp4. HEK 293T cells were seeded into 24-well plates, and X-tremegene HP (ROCHE) was used to transfect the plasmid. MiR-29a-5p mimics were transfected into HEK 293T cells with wild-type and mutant Gsk3b 3’UTR-carrying luciferase. A Passive Lysis Buffer was added to the well to lyse cells. The Luciferase Assay Reagent (Promega) was used to test firefly luminescence. Stop & Glo Reagent was used to test Renilla luminescence. The effect of miR-29a-5p on the activity of the Gsk3b luciferase reporter gene was observed. In addition, dual-luciferase activity technology was used to clarify the relationship between miR-29a-5p and Aqp4 as the above method.

### 2.15. Statistical analysis

Statistical analyses were conducted using SPSS 26.0. All measurement data were expressed as mean ± standard deviation. The Shapiro-Wilk test was used to determine whether econometric data were normally distributed. One-way analyses of variance followed by the Least Significant Difference were performed to compare among three or more groups. The level of significance was set at 0.05.

## 3. Results

### 3.1. Increased miR-29a-5p alleviated astrocyte injury and regulated the expression of A1 and A2 phenotypes of reactive astrocytes in the OGD/R model

In OGD/R-treated astrocytes, there was decreased expression of miR-29a-5p at 24 hours. The miR-29a-5p mimics increased the expression of miR-29a-5p in astrocytes following OGD/R, while miR-29a-5p inhibitors decreased the expression of miR-29a-5p (Figure 1A). The miR-29a-5p mimics alleviated the astrocyte damage caused by CCK-8 (p<0.001), inhibited LDH release from astrocytes (p=0.001) in OGD/R-treated astrocytes. The miR-29a-5p inhibitors worsened cell damage (P<0.01) induced by OGD/R (Figure 1B and 1C). The dynamic expression of the A1 reactive astrocyte markers (C3, Fkbp5, Serping1, Ggta1, Gbp2) and A2 reactive astrocyte markers (S100a10, Ptx3, S1pr3, Emp1, Cd109) following OGD at 0, 3, 6, 12, and 24 hours after reoxygenation was shown in supplementary Figure S1. The expression of the A1 markers of C3, Fkbp5, and Serping1 was decreased (p<0.001), while the expression of the A2 markers of S100a10, Ptx3, and Emp1 was increased (p<0.01) after intervention with miR-29a-5p mimics compared to the OGD/R-treated group in astrocytes (Figure 1D and 1E). It seems that upregulated miR-29a-5p can alleviate astrocyte injury, suppress A1-type astrocyte expression, and promote the activation of A2-type astrocytes after OGD/R.

**Figure 1.**
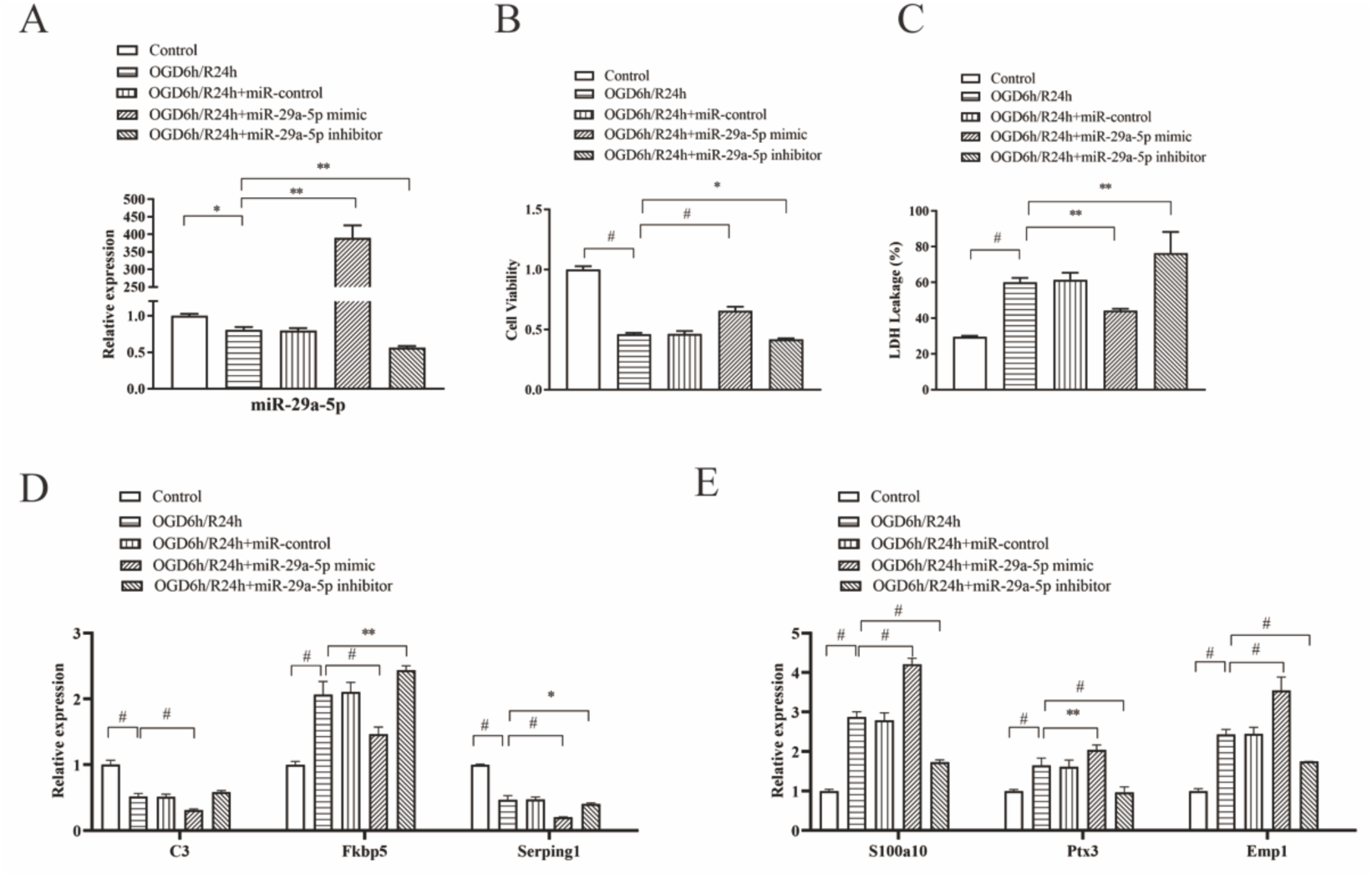
Elevating miR-29a-5p levels was protective in ischemia-reperfusion injury in the OGD/R astrocyte model. **A**, miR-29a-5p mimics increased the expression of miR-29a-5p in astrocytes following OGD/R. **B**, miR-29a-5p mimics alleviated cell damage shown by CCK-8 assay. **C**, miR-29a-5p mimics inhibited LDH release from astrocytes. **D and E**, miR-29a-5p mimics decreased the expression of neurotoxic A1 reactive astrocyte markers (C3, Fkbp5, and Serping1) while increasing the expression of neuroprotective A2 reactive astrocyte markers (S100a10, Ptx3, and Emp1). Each sample was tested in triplicate. *P<0.05; **P<0.01; #P<0.001.

### 3.2. Increased miR-29a-5p regulated the expression of predicted target genes for miR-29a-5p in the OGD/R model

Predicted target genes for miR-29a-5p were identified and were analyzed for biological function. The enriched KEGG pathways of the predicted target gene were revealed by DAVID Bioinformatics Resources. (Supplementary Figure S2A). Then, we selected 245 predicted targeted genes in the top ten KEGG pathways to establish the protein-protein interaction network (Supplementary Figure S2B). The top twenty hub genes were identified (Figure 2A and Supplementary Table S1). We found twelve hub genes with significantly differential expression in the astrocyte OGD/R model.

**Figure 2.**
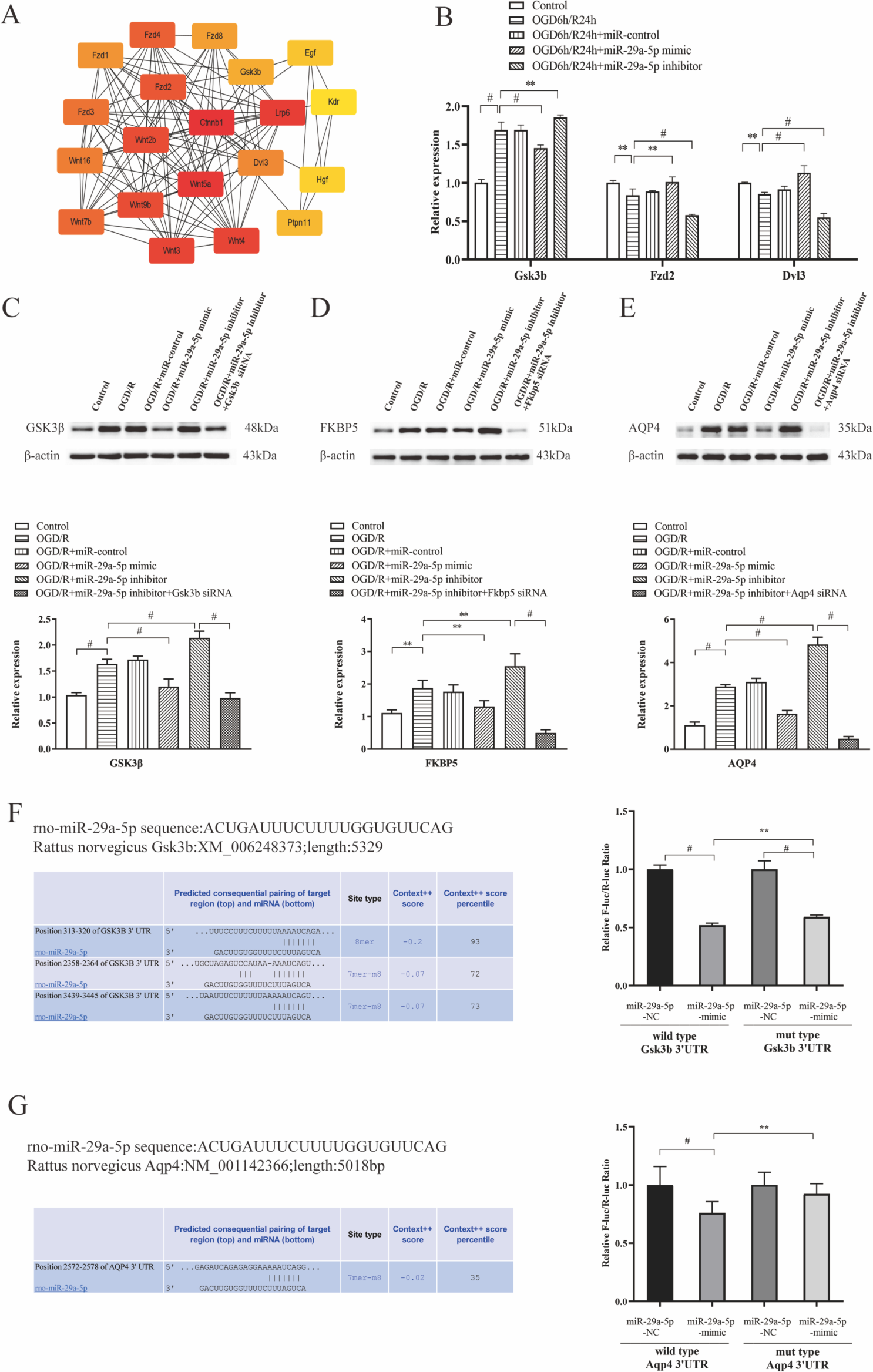
miR-29a-5p regulated the expression of target genes in the OGD/R astrocyte model. **A**, the top predicted twenty hub genes for miR-29a-5p were identified by bioinformatics analysis. **B**, miR-29a-5p mimics suppressed the upregulated expression of Gsk3b and increased the downregulated expression of Fzd2 and Dvl3 shown by RT-PCR. **C, D, and E**, Western blot analysis showed miR-29a-5p mimics significantly suppressed the upregulated expression of GSK3β, FKBP5, and AQP4. **F and G**, Gsk3b and Aqp4 are the direct target genes for miR-29a-5p by a dual-luciferase reporter assay. Each sample was tested in triplicate. **P<0.01; #P<0.001.

Five hub genes were significantly upregulated 24 hours after OGD/R, including Wnt family member (Wnt) 4, Wnt2b, frizzled class receptor (Fzd) 4, Fzd8, and Gsk3b.

Seven hub genes were significantly downregulated 24 hours after OGD/R, including β-catenin, Wnt5a, lipoprotein receptor-related protein 6 (Lpr6), Fzd2, Fzd3, disheveled segment polarity protein (Dvl) 3, and hepatocyte growth factor (Supplementary Figure S3).

MiR-29a-5p mimics suppressed the expression of Gsk3b and increased the expression of Fzd2 and Dvl3 among the twelve predicted hub genes in OGD/R-treated astrocytes (p<0.01) (Figure 2B). The other nine predicted hub genes were not significantly regulated after intervention with miR-29a-5p mimics or inhibitors.

### 3.3. MiR-29a-5p regulated the expression of GSK3β, FKBP5, and AQP4 in the OGD/R model

MiR-29a-5p mimics suppressed the upregulated expression of GSK3β in OGD/R-treated astrocytes shown by Western blot analysis (p<0.001). Silencing the Gsk3b gene weakened the effect of the miR-29a-5p inhibitors on the expression of GSK3β in the astrocyte OGD/R model (Figure 2C). MiR-29a-5p mimics suppressed the upregulated expression of FKBP5 in OGD/R-treated astrocytes (p<0.01). Silencing the Fkbp5 gene weakened the effect of the miR-29a-5p inhibitors on the expression of FKBP5 in the astrocyte OGD/R model (Figure 2D). MiR-29a-5p mimics suppressed the upregulated expression of AQP4 in OGD/R-treated astrocytes (p<0.001). Silencing the Aqp4 gene weakened the effect of the miR-29a-5p inhibitors on the expression of AQP4 in the astrocyte OGD/R model (Figure 2E).

### 3.4. Gsk3b and Aqp4 are the direct target gene for miR-29a-5p

The inhibitory effect of miR-29a-5p mimics on the target gene Gsk3b 3’UTR and Aqp4 3’UTR were detected using the dual-luciferase reporter assay. Site-specific mutagenesis technology determined the site of interaction between miR-29a-5p and Gsk3b 3’UTR. The miR-29a-5p mimics significantly inhibited the luciferase activity expressed in tandem with the wild-type Gsk3b 3’UTR (P<0.001) and partially inhibited the luciferase activity expressed in tandem with the mutant Gsk3b 3’UTR; however, the inhibitory ability was partially weakened. These results indicated that Gsk3b is a direct target gene for miR-29a-5p (Figure 2F).

Site-specific mutagenesis technology determined the site of interaction between miR-29a-5p and Aqp4 3’UTR. The miR-29a-5p mimics significantly inhibited the luciferase activity expressed in tandem with the wild-type Aqp4 3’UTR (P<0.001). These results indicated that Aqp4 is a direct target gene for miR-29a-5p (Figure 2G).

### 3.5. Increased miR-29a-5p regulated the expression of miR-29a-5p, A1 and A2 astrocyte markers in the reperfusion-induced HT model

The levels of miR-29a-5p were decreased in both the peri-infarction tissue and whole blood following reperfusion at 0 and 6 hours in the rat MCAO model (Figure 3A). Intravenous administration of miR-29a-5p agomir before reperfusion increased the expression of miR-29a-5p in both the peri-infarction and whole blood compared with the ischemia-reperfusion group (P<0.001) (Figure 3B).

**Figure 3.**
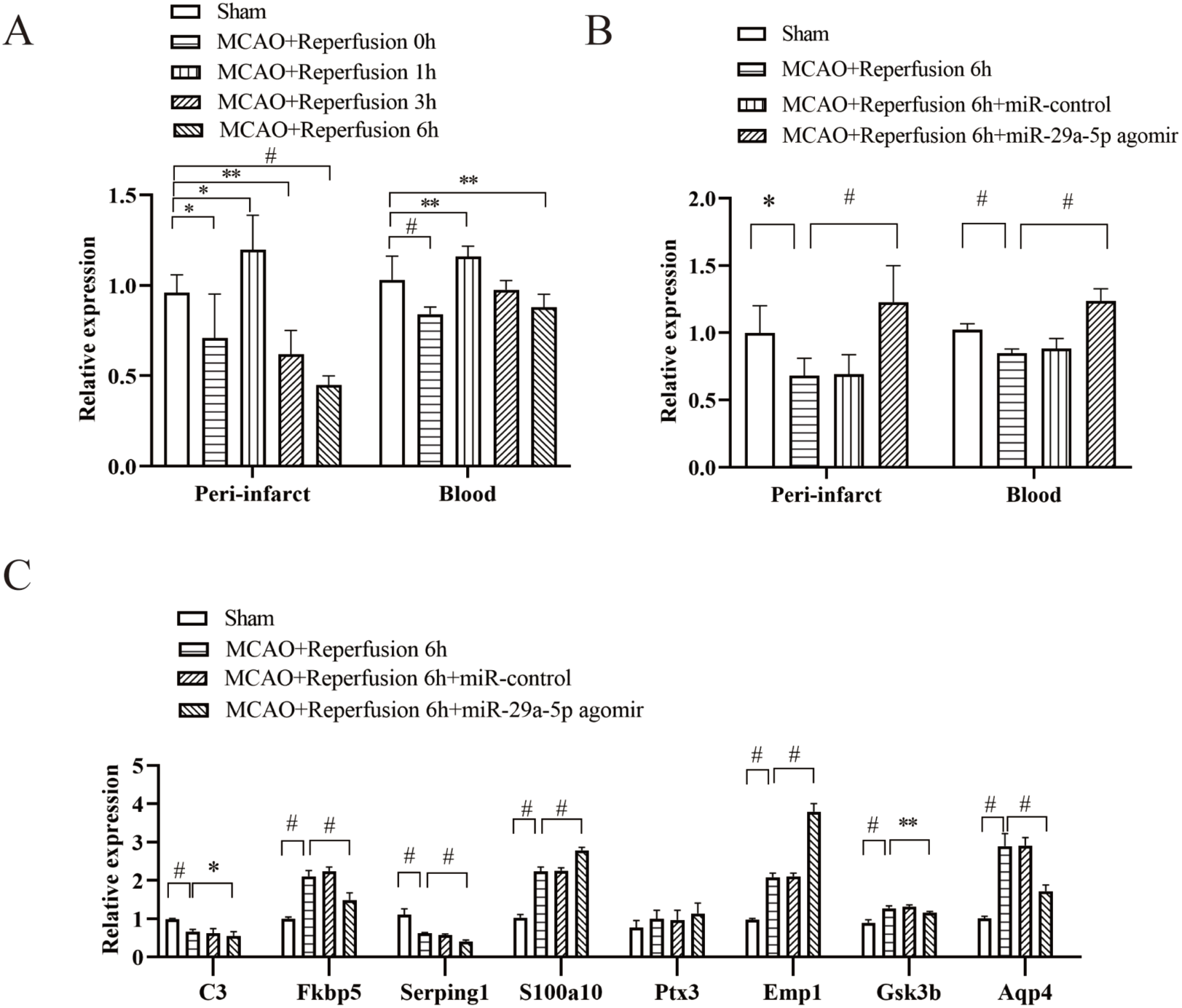
Elevating miR-29a-5p levels regulated the expression of miR-29a-5p, predicted target genes, and A1/A2 astrocyte markers in the reperfusion-induced HT model. **A,** miR-29a-5p expression decreased in the peri-infarction tissue and blood of the MCAO model (n=6). **B,** miR-29a-5p agomir treatment before reperfusion reduced the miR-29a-5p expression in the peri-infarction tissue and blood of the MCAO model (n=6). **C,** miR-29a-5p agomir treatment suppressed the mRNA expression of A1 markers (C3, Fkbp5, and Serping1), Gsk3b, and Aqp4, while enhanced A2 markers (S100a10 and Emp1) in the peri-infarction tissue (n=6). *P<0.05; **P<0.01; #P<0.001.

The mRNA expression of the A1 astrocyte markers (C3, Fkbp5, and Serping1) was decreased, while the expression of the A2 astrocyte markers (S100a10 and Emp1) was increased in the peri-infarction tissue after intervention with miR-29a-5p agomir (Figure 3C).

### 3.6. Increased miR-29a-5p reduced infarct volume, HT, BBB breakdown, and improved functional outcome

MiR-29a-5p agomir treatment before reperfusion reduced cerebral infarct volume (TTC staining, 15.7% versus 34.7%, p<0.001) (Figure 4A) and reduced HT (hemoglobin content index, 1.4 versus 1.9, p=0.001) (Figure 4B), compared with the ischemia-reperfusion group in the rat MCAO model. There was less Evans blue extravasation after miR-29a-5p agomir treatment (1.9 versus 3.7, p<0.001), indicating the protection of BBB by increased the level of miR-29a-5p (Figure 4C). Rats treated with miR-29a-5p agomir showed better neurological scores following reperfusion at 6 hours (1.1 versus 2.2, p<0.001).

**Figure 4.**
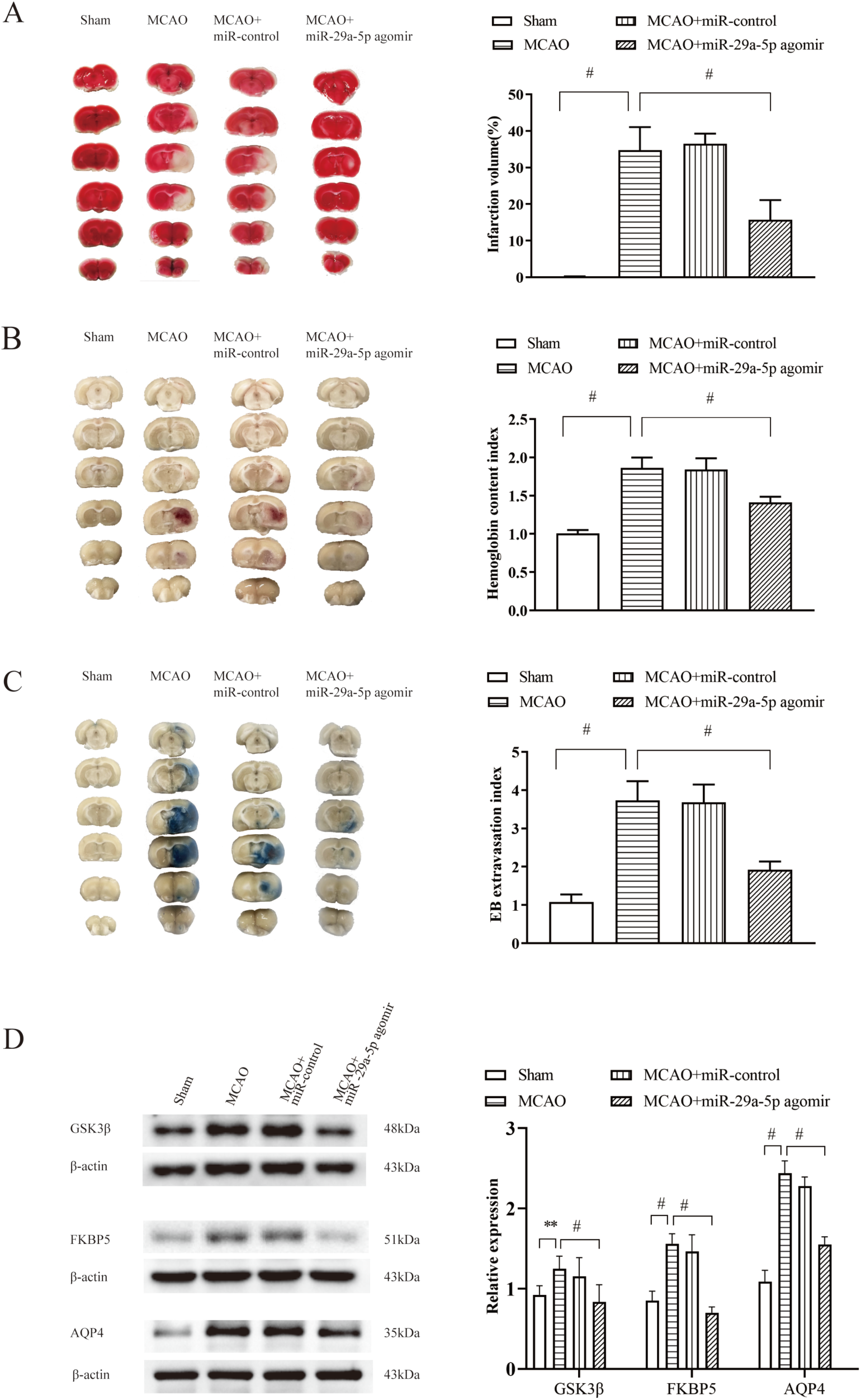
MiR-29a-5p agomir reduces infarct volume, reperfusion-induced hemorrhage, and BBB disruption and improves neurological outcomes via regulating GSK3β, FKBP5, and AQP4. **A**, TTC staining of brain slices and quantification for infarct volume (n=6). **B**, Hemoglobin content index is determined by spectrophotometric hemoglobin assay (n=6). **C**, BBB permeability in the ischemic hemisphere was shown by Evans blue staining (n=6). **D**, Western blot shows the expression of GSK3β, FKBP5, and AQP4 in the peri-infarction tissue (n=6). **P<0.01; #P<=0.001.

### 3.7. Increased miR-29a-5p regulated the expression of GSK3β, FKBP5, and AQP4 in the reperfusion-induced HT model

The mRNA expression of Gsk3b and Aqp4 as the direct target gene for miR-29a-5p was decreased in the peri-infarction tissue after intervention with miR-29a-5p agomir (Figure 3C). In addition, miR-29a-5p agomir treatment suppressed the increasing expression of GSK3β, FKBP5, and AQP4 in the peri-infarction tissue compared with the ischemia-reperfusion group (p<0.001) shown by Western blot analysis (Figure 4D).

## 4. Discussion

Our study provides novel evidence supporting the relationship between astrocytic miR-29a-5p and HT in ischemic stroke using an in vitro astrocyte OGD/R model and a reperfusion-induced HT rat model. The miR-29 family is significantly downregulated in cerebral ischemia-reperfusion injury, influencing cellular responses to ischemic damage [15–18]. Specifically, miR-29a-3p has been observed to decrease in the CA1 region in a rat forebrain ischemia model, where it targets the pro-apoptotic protein PUMA, thus mitigating ischemic injury in astrocytes and neurons in vitro [15, 30]. Additional research has shown that miR-29a-3p overexpression protects neuronal cells from OGD/R injury by downregulating Rock1 and modulating TNFRSF1A and NF-κB signal pathways [31,32]. MiR-29a-5p was found to be reduced in the plasma of stroke patients, correlating with the severity of brain injury and clinical outcomes. Knocking out miR-29a resulted in the increased M1 microglia polarization, thereby increasing infarct volume in a rat stroke model [19]. Collectively, these findings align with our results, suggesting that miR-29a-5p overexpression may reduce cerebral infarction volume, mitigate BBB disruption and HT, and improve functional outcomes in ischemic stroke models.

Our findings show that miR-29a-5p overexpression can attenuate astrocyte injury, decrease expression of neurotoxic A1 markers (C3, Fkbp5, and Serping1), and increase neuroprotective A2 markers (S100a10 and Emp1) following OGD/R and mechanical reperfusion after MCAO. These results suggest that miR-29a-5p promotes a neuroprotective environment that supports neuronal survival and tissue repair. In ischemic stroke contexts, reactive astrocytes can adopt either an A1 phenotype, associated with neurotoxicity and increased pro-inflammatory cytokine release, or an A2 phenotype, linked with neuroprotection and the secretion of neurotrophic and anti-inflammatory factors [23,24]. Increased S100a10 expression, a hallmark of A2 astrocytes, has shown protective effects in cerebral ischemia-reperfusion injury [33]. Moreover, the astrocyte-specific Kruppel-like factor 4 has been shown to suppress A1 activation and promote A2 polarization post-ischemia, primarily through NF-κB pathway modulation [24]. Research underscores that promoting the A2 phenotype can enhance post-stroke recovery by supporting neuronal survival and reducing inflammation [22,33]. Our results on the ability of miR-29a-5p to modulate astrocyte phenotypes add to this growing body of evidence, suggesting potential therapeutic strategies for reducing HT and improving post-stroke outcomes.

This study also reveals that increasing miR-29a-5p levels effectively downregulates its direct and predicted target gene in vitro and in vivo, illustrating its protective role in astrocytes. Our results indicate that miR-29a-5p exerts this neuroprotection by regulating genes such as Gsk3b, Aqp4, and Fkbp5. Prior studies suggest that Gsk3b inhibition can protect the BBB and alleviate ischemia-reperfusion injury in ischemic stroke models [34]. Wnt-3a protein administration leads to Gsk3b dephosphorylation, reducing cerebral infarction and neuronal apoptosis through the Foxm1 pathway [35]. Aqp4, an astrocyte endfeet marker, is upregulated in ischemic tissue following BBB disruption [36]. Notably, miR-29b overexpression has been shown to reduce BBB disruption in ischemic stroke by downregulating Aqp4 [16].

Nicotinamide adenine dinucleotide phosphate oxidase inhibitor has been found to decrease brain edema and HT by downregulating Aqp4 in the reperfusion-induced HT model [20]. We also identified Fkbp5 as a predicted target gene for miR-29a-5p. In a previous study, Fkbp5 was upregulated in cerebral ischemia-reperfusion injury and regulated autophagy via the AKT/FOXO3 signaling pathway [37]. In our study, we observed increased Fkbp5 levels in OGD/R-treated astrocytes and peri-infarction tissue of the HT model, which were reduced by miR-29a-5p overexpression.

Our study has certain limitations. We did not evaluate the long-term effects of miR-29a-5p modulation on astrocyte function and neuroprotection, nor did we investigate potential side effects associated with miR-29a-5p modulation. Further research is warranted to clarify the direct interaction between miR-29a-5p and the predicted target gene of Fkbp5.

## 5. Conclusions

Our study underscores the protective role of miR-29a-5p in astrocytes under ischemia-reperfusion injury conditions. By targeting Gsk3b, Aqp4, and Fkbp5, miR-29a-5p effectively reduces astrocyte injury and fosters a neuroprotective environment. MiR-29a-5p could serve as a therapeutic target to minimize HT and improve outcomes following mechanical reperfusion in AIS.

## Author contribution statement

Z-SS participated in the design of the present study. All authors participated in the interpretation and collection of the data. C-LL, J-KZ and Z-SS wrote the initial manuscript. Z-SS revised the manuscript. All authors critically reviewed and edited the manuscript and approved the final version.

## Ethics statements

The study was reviewed and approved by the Institutional Animal Care and Use Committee of Sun Yat-sen Memorial Hospital, Sun Yat-sen University.

